# MetaPointFinder: Detection of Antimicrobial Resistance-Conferring Point Mutations (ARMs) from Metagenomic Reads

**DOI:** 10.64898/2025.12.04.691125

**Authors:** Aldert L. Zomer, Sam Nooij, Cailean Carter, Alison E. Mather

## Abstract

**Background:** Many clinically important antimicrobial resistance (AMR) phenotypes such as fluoroquinolones and rifamycins are driven by antimicrobial resistance-conferring mutations (ARMs) in conserved chromosomal loci (e.g., *gyrA, parC, rpoB*). Resistome profiling by metagenomics sequencing is often proposed as an ideal AMR surveillance tool as it is organism-agnostic, but currently, existing metagenomic AMR surveillance pipelines are only able to identify acquired AMR genes but not point mutations associated with AMR. This is a serious gap in metagenomics-based AMR surveillance, as the true extent of AMR may be underestimated.

**Methods:** We developed MetaPointFinder (v1), a read-based method that can process both long and short metagenomic reads. The tool identifies resistance-determining regions in reads using DIAMOND (translated protein) and KMA (nucleotide), and classifies known resistant versus wild-type variants by aligning the sequences using pwalign and assessing the observed mutations based on known resistance mutations in the AMRFinderPlus database. The tool outputs ARMs per read, per gene and per antibiotic class.

**Results:** In proof-of-concept analyses, MetaPointFinder identified known AMR-associated mutations and quantified resistant/susceptible read counts and ratios from metagenomic samples with corroborating phenotypic resistance data when available. We benchmark our tool using simulated reads from DNA and from reverse-translated protein references with read lengths of 100–5000 bp and error rates between 0 and 30%, simulating both Illumina and Nanopore error rates. We show that MetaPointFinder outperforms any available method for detection of ARMs in metagenomics data.

**Conclusion:** MetaPointFinder complements gene-centric resistome profiling by capturing chromosomal mutation-based AMR directly from metagenomes.

**Impact Statement:** Many clinically important antimicrobial resistance (AMR) phenotypes such as fluoroquinolones and rifamycins are driven by antimicrobial resistance-conferring mutations (ARMs) in conserved chromosomal loci (e.g., *gyrA, parC, rpoB*). Resistome profiling by metagenomic sequencing is ideal for AMR surveillance, but existing pipelines detect only acquired genes and ignore point mutations. MetaPointFinder bridges this gap. The tool accurately identifies reads carrying resistance-determining regions using DIAMOND (protein) and KMA (DNA), aligns them, and classifies resistant versus wild-type variants using known mutations from the AMRFinderPlus database.

## Introduction

Antimicrobial resistance (AMR) is a threat to human and animal health globally, and the rapid and accurate quantification of AMR is critical to any strategy to reduce its burden. Generation and availability of bacterial sequence data and lower costs compared to phenotypic antibiotic susceptibility testing makes genotypic prediction of AMR increasingly common. Many tools exist that facilitate the prediction of AMR from genome sequences such as ResFinder [1], AMRFinderPlus [2] or RGI [3]. Most available tools predict resistance from both acquired antimicrobial resistance genes (ARGs), and from antimicrobial resistance mutations (ARMs) in resistance-determining regions (RDRs) which are often found in essential genes. The identification of genotypic resistance does not necessarily equate to phenotypic resistance, but concordance between phenotype and genotype is generally fairly high [4–7]. Whole genome-based AMR surveillance requires culturing of isolates and this restricts the assessment of AMR to specific targeted bacteria. An organism-agnostic alternative which does not require culturing is metagenomic sequencing, where estimation of the resistome can be obtained either by mapping reads to ARG databases or by assembling metagenomic reads into contigs and looking for ARGs in the assemblies [8–10].

Metagenomic sequencing has transformed ARG profiling, as it offers a more complete overview of the types and diversity of ARGs within samples. It has been applied in a variety of different settings across the One Health continuum, including surveillance of AMR in wastewater [11, 12], food [13, 14], humans [15, 16] and animals [17]. Typically, the methods used only evaluate the abundance of ARGs and are not able to identify ARMs because identification of single nucleotide polymorphisms (SNPs) in metagenomic data is not trivial because it is difficult to validate performance. The tool Mumame, developed to fill this gap, identifies mutations but displayed highly variable quantitative estimates, even at high sequencing depths [18]. This is a serious gap in metagenomics-based AMR surveillance, as the total AMR of a sample is likely to be underestimated if ARMs cannot be reliably identified. ARMs underpin resistance to several frontline classes of antibiotics such as fluoroquinolones, rifamycins, and folate-pathway inhibitors, yet remain undetected in standard AMR pipelines. Although tools such as ResFinder and AMRFinderPlus, implemented in many whole genome sequencing and AMR surveillance pipelines, are highly successful for isolate genome assemblies, they cannot be run in a species-agnostic manner, necessary for metagenomic samples where multiple bacterial species are present and where minor variant mutations are lost in non-haplotype aware assemblies. There remains a practical gap for robust, read-based identification and quantification of known resistance-conferring substitutions directly from metagenomes.

We present MetaPointFinder (v1), a species-agnostic, read-based approach that detects known SNPs and amino-acid substitutions in resistance-determining regions (RDRs) using translated and nucleotide alignment and reports resistant/susceptible/unknown read counts. We show the detection of substitutions in the quinolone resistance-determining region of DNA gyrase subunit A gene GyrA (e.g., S83L, D87N), where association with fluoroquinolone resistance is well established and compare with available phenotypic data, we show the detection of SNPs and amino-acid substitutions and evaluate ARM counts in various public datasets. Finally, we also evaluate Mumame [18], an earlier mutation-mapping approach using Usearch [19], which we compare our tool against in a benchmark generated from synthetic reads containing either known wildtype or resistance-causing mutations at different read lengths and error rates.

## Materials and Methods

### Reference targets and databases

All resistance-determining regions (RDRs) were extracted from the AMRFinderPlus [2] catalogue (genes with known SNPs or proteins with known substitutions) and stored as nucleotide and protein entries with metadata (gene, accession, position in reference sequence, mutation(s), antimicrobial class) (accessed 17^th^ of July 2025).

### Alignment strategy

FASTQ reads were six-frame translated and aligned with DIAMOND blastx v 2.1.15 [20] using the flags “-F 15 --identity <70-85> --range-culling -k1 --range-cover 5 --iterate --masking 0 --outfmt 6 qseqid sseqid pident length mismatch gapopen qstart qend sstart send evalue bitscore full_sseq qseq_translated” followed by precise alignment and indexing every position using pwalign [21] with the following parameters: “BLOSUM62; gap open −10, gap extend −0.5; local–global scoring”. Reads were also aligned to RDR DNA sequences with KMA 1.4.15 [22] using the following command line parameters “-bcNano -hmm -ont -nc -na -1t1 -mrs <0.5-0.85>“; followed by precise alignment using pwalign [21] with the following parameters: “match = 2, mismatch = −5, baseOnly = FALSE, gapOpening = −10, gapExtension = −0.5, local–global scoring”.

### Variant calling and classification

From the alignments, MetaPointFinder extracts nucleotides or amino acids at canonical positions and classifies each read as resistant or wildtype using the curated mutation table obtained from AMRFinderPlus [2]. The tool then lists and counts the ARMs and non-ARMs detected per read per class; to note, a read can have multiple ARMs and multiple RDRs for different classes. Ambiguities, such as non-covered regions, insertions or deletions (indels), unreported substitutions/mutations and unknown amino acids (X), are detected and scored as “unknown”. Resistant, wild-type and unknown mutations are reported per read, per locus and per antibiotic class.

### Synthetic benchmark design

To enable head-to-head comparisons with Mumame, the only known previously published read-centric AMR metagenomic analysis tool, we first randomly selected three known mutations from the MetaPointFinder database from each protein and gene in the database and generated protein and gene sequences with and without the mutations. Following, we reverse-translated the protein references using EMBOSS backtranseq (https://emboss.sourceforge.net/ v6.6.0) with the *E. coli* codon usage table to DNA sequences. Random nucleotides were padded on both ends of the inferred DNA sequences up to 6 kb and reads where simulated from these 6 kb fragments using wgsim 0.3.1-r13 (https://github.com/lh3/wgsim) at read lengths 100, 200, 1000, and 5000 bp with error rates between 0 and 30% of which 15% of polymorphisms are indels. Between *1*.*0 × 10*^*8*^ and *1*.*0 × 10*^*9*^ bases of sequence data were generated to ensure that approximately *2*.*0 × 10*^*4*^ reads contained an RDR per FASTQ file, with roughly *1*.*0 × 10*^*4*^ reads representing the wild-type variant and *1*.*0 × 10*^*4*^ reads representing the mutated variant. For DNA benchmarks, nucleotide sequences were padded up to 6 kb with random sequences and processed as described. The benchmark repository and scripts are available here: https://github.com/aldertzomer/metapointfinder/tree/main/benchmark and can be rerun from a functional MetaPointFinder installation. Outputs of the benchmark code include the numbers of resistant and wild-type genes detected, and correctly and incorrectly mapped reads per gene. MetaPointFinder was run with the following settings “--identity 50” for the benchmarks. Mumame database indexing was performed using “mumame_build -m snps.txt -i protein_fasta_protein_variant_model.fasta” which relies on the latest CARD database (v4.0.1) [3] and provides the necessary files and the analysis was run on the synthetic benchmark FASTQ files using default settings.

### Real-data re-analyses

To evaluate the performance of MetaPointFinder against real-world datasets, we re-analysed five datasets: (i) An Illumina sequencing dataset associated with prolonged selection for resistance after antimicrobial treatment of chickens, focusing on fluoroquinolone and β-lactam resistance signals where genotypic–phenotypic discrepancies have been described [23], obtained from ENA SRA PRJEB73721. (ii) An Illumina sequencing dataset of tuberculosis (TB) patients treated with rifampicin [24], ENA SRA PRJNA1002577. (iii) An Illumina sequencing dataset of sewage samples by Munk et all [12] (see Supplemental Data 1 for all accessions, only read files with more than 5 gbase of sequencing data were included). (iv) A *gyrA*/*parC* amplicon sequencing dataset where no difference was observed between the encoded GyrA/ParC resistance substitutions in Swedish and Indian wastewater treatment plants (WWTPs), now analysed without discarding a majority of reads [25], ENA SRA PRJNA239415. (v) The abundance of ARGs and ARMs from a set of 1228 long-read metagenomes obtained from SRA with more than 1 gbase of sequence data per readfile. In all cases, reads were downloaded from SRA and analysed without any pre-processing. Resistance fractions were determined by dividing resistant RDR counts by the sum of the wildtype and resistant RDR counts. Relative RDR abundances were determined by dividing the number of resistant RDR counts by the total read counts in file, then multiplying by 1,000,000 to scale the relative abundance to 1,000,000 reads. Heatmaps were generated from RDR fractions or scaled relative abundances using pheatmap [26] using Euclidian distance clustering. RDRs with less than 10 reads mapped were excluded from the analysis. Barcharts were generated in Excel or GraphPad Prism v10 with whiskers displaying 95% confidence intervals.

### Statistical testing

RDR counts or percentages were compared using a t-test or with a repeated measurements ANOVA with Benjamini Hochberg correction for multiple testing in GraphPad Prism v10. Significance (p<0.05) is depicted by “*” in the graphs. Correlation was performed using by Pearson correlation on (sequencing depth adjusted) readcounts, log(x + 1) transformed readcounts or by Spearman Rank correlation on sequencing depth adjusted read counts with and without zero counts excluded to avoid spurious correlation results.

### Implementation

MetaPointFinder v1 is implemented in Python 3 (v.3.11.9 was used) and utilises DIAMOND and KMA for detection of RDR-containing reads and performs precise alignments in R v4.4.0 using pwalign v1.2 in Bioconductor 3.20 and Parallel for multiprocessing speedup. It detects ARM containing reads, parses alignments for precise determination and outputs tabular summaries (*.class.prot.summary.txt; *.class.dna.summary.txt) using Biostrings 2.74.1 from Bioconductor 3.20. The pipeline accepts (gzipped) FASTQ (short- or long-read) and is species-agnostic. Code is available here: https://github.com/aldertzomer/metapointfinder. Conda and Docker packages are available. A web-based implementation is available here: https://klif.uu.nl/metapointfinder/.

## Results

### Simulated benchmark performance

Gold standard non-simulated sequencing datasets are not available for ARM data, therefore we simulated FASTQ-formatted read files from the MetaPointFinder database which contains 260 proteins and 25 DNA regions. Approximately 20,000 reads out of 20,000 – 1,000,000 reads (number of reads depends on the simulated read length) contain an actual RDR per FASTQ file, with approximately 10,000 reads containing a wildtype variant and 10,000 reads containing a mutated variant. Reads were both generated from RDRs with amino-acid substitutions and from RDRs with DNA mutations, although the latter could not be tested by Mumame as all attempts to run Mumame using a DNA mutation database failed.

On all simulated datasets, MetaPointFinder recovered 95-100% of all known resistant and wildtype RDRs up to an introduced error rate of 0.2 (Fig 1A, blue lines). Mumame detected only 40-55% of the RDRs (Figure 1B, blue lines) at error rates of 0 – 0.03 and then the detection of RDRs drops. A lower error rate (10-1000 fold lower) was observed for MetaPointFinder when comparing with Mumame (Figure 1D vs 1E, orange lines). MetaPointFinder was less sensitive to sequence divergence caused by higher error rates: it still detected genes and mutations with error rates up to 30% (Fig 1A vs 1B). However, the number of false positives increased as well, likely caused by changes to the actual ARM (at an error rate of 0.3, almost 1 in 3 bases is changed from the reference). Practically no RDRs were detected with Mumame at error rates of 0.2 or higher and as a consequence the number of false positives was also 0 (Fig 1B, 1E).

**Figure 1.**
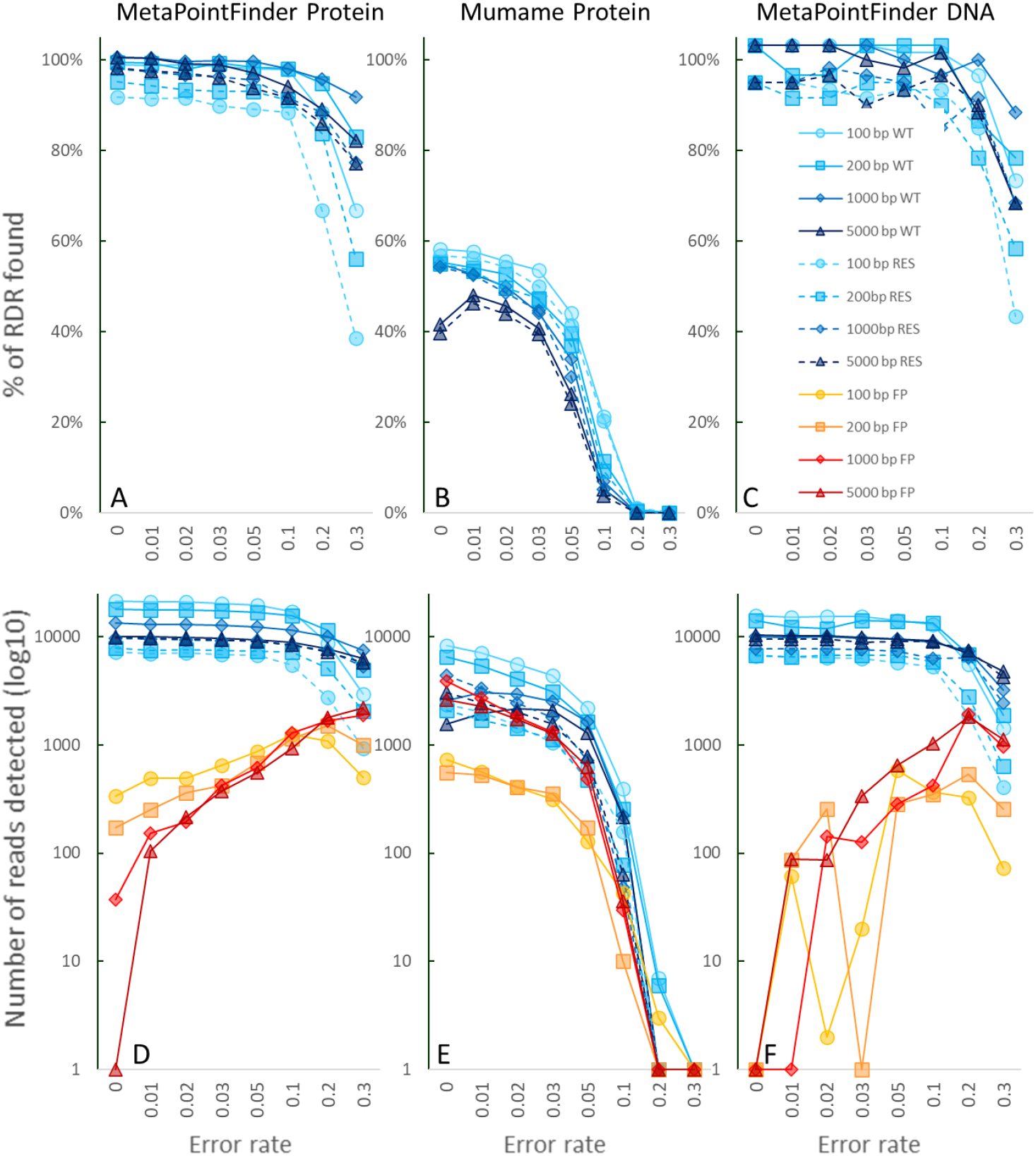
Benchmark data from synthetic read data generated from proteins and genes with known wild-type or resistance mutations for various read lengths and error rates causing divergence from actual sequence (0-0.3 = 0%-30%). Connected lines: Data from reads originating from wildtype RDRs (WT). Dotted lines: Data from reads originating from resistant RDRs (RES). In blue shades: correctly predicted. In orange-red shades: false positive (FP: wildtype reads predicted as resistant). A: MetaPointFinder amino-acid performance. B: MUMAME amino-acid performance C: MetaPointFinder DNA performance. D: MetaPointFinder RDR amino-acid performance. E: MUMAME RDR amino-acid performance. Row F: MetaPointFinder RDR DNA performance.

### Illumina fluoroquinolone and beta-lactam resistome data

To evaluate the effectiveness of MetaPointFinder on Illumina sequencing data, we investigated a longitudinally sampled dataset from Swinkels et al. [23] of chickens treated with enrofloxacin, amoxicillin and doxycycline. Here, phenotypic fluoroquinolone resistance in isolated *E. coli* from samples was clearly observed after enrofloxacin treatment for all timepoints, with all isolates showing resistance to enrofloxacin and carrying resistance causing mutations in the gene coding for GyrA. The authors reported that Illumina metagenomic analysis of the faecal samples did not show an appreciable increase in the fluoroquinolone resistome; the metagenomes were searched for acquired ARGs. We analysed the Illumina data and show that a statistically significant relative increase in fluoroquinolone ARMs could be observed in the enrofloxacin treated flocks (Figure 2A). A statistically significant relative increase in beta-lactam ARMs was also observed in the amoxycillin treated flocks (figure 2B) in timepoint 1, similar as what was observed for the phenotypic and ARG resistome data in the original study. No increase was observed for ARMs for tetracyclines as few ARMs were detected (not shown). Interestingly, the relative increase in quinolone resistance seems to be mostly caused by disappearance of the strains not carrying a resistance causing mutation (Figure 2C) for the enrofloxacin treated flocks.

**Figure 2.**
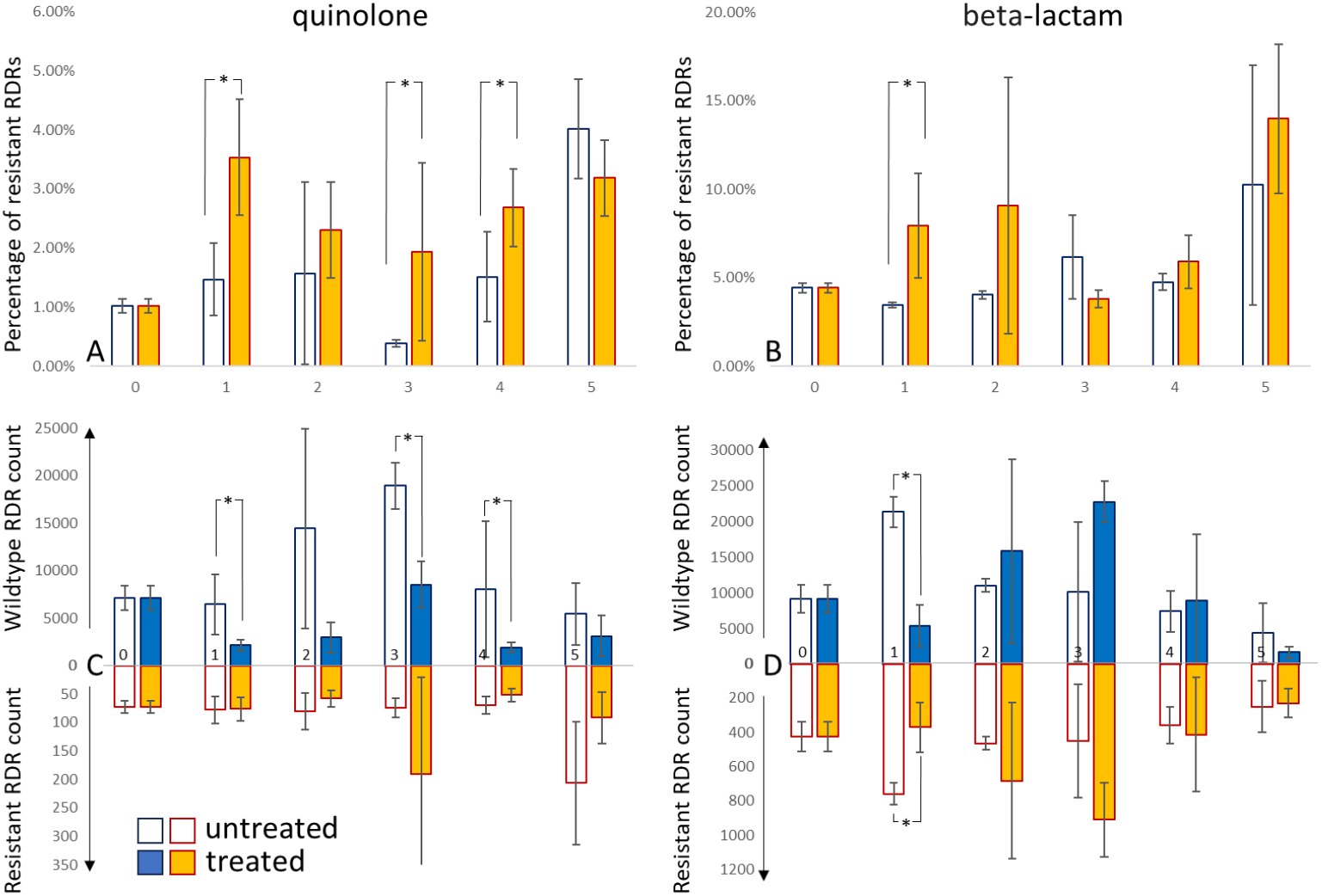
Percentage resistance and read counts of reads predicted to carry a wildtype base(s) or resistance mutation, based on re-analysis of the metagenomics sequence data from Swinkels et al [23]. Left (AC): Quinolone resistance in enrofloxacin treated flocks vs controls. Right (BD): Beta-lactam resistance in amoxicillin treated flocks. Top (AB). Y-axis: % of reads predicted to carry a resistance mutation. X-axis: timepoint. Bottom (BD): Y-axis: number of reads predicted to carry a wildtype or resistance mutation. X-axis: timepoint. Significant differences are marked by * (p<0.05, t-test).

### Illumina rifamycin resistome data

To evaluate the performance of MetaPointFinder on detecting RpoB amino acid substitutions in metagenomes, we re-analysed sequence data of Diallo et al [24]. Here, the gut microbiome of tuberculosis (TB) patients was sequenced using Illumina sequencing at timepoint 0 (before treatment), at 2 and 6 months (during treatment with rifampicin) and at 15 months (9 months after treatment). Exposure to rifampicin generally results in mutations in RpoB in many bacterial species [27]. In the original study, the authors showed that the gut microbiome of TB patients during changes during treatment and the altered microbial community in the gut environment persisted for at least 9 months after treatment completion. The authors did not discuss the resistome for us to make comparisons with. Using MetaPointFinder, we show that resistance to rifamycins was higher in treated patients during treatment, but that the rifamycin resistance in the gut microbiome after treatment was no longer different to the before treatment sample (Figure 3). RpoB mutations come with a fitness cost and without antimicrobial pressure, resistant strains are lost from the microbiome [27].

**Figure 3.**
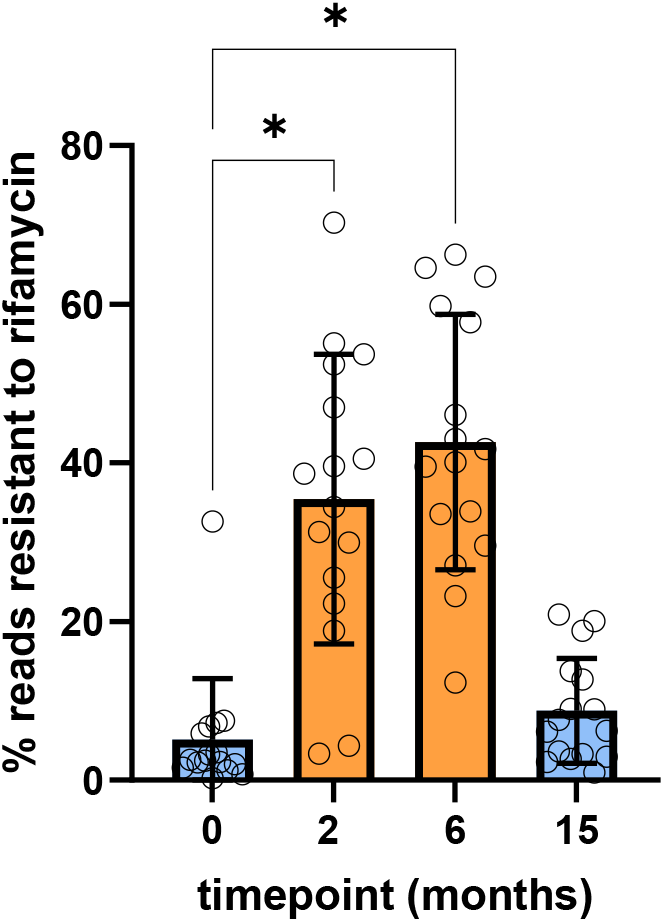
Percentage of reads mapped to rifamycin RDRs resistant to rifamycin as compared to wildtype reads in 4 timepoints from Diallo et al [24]. In orange: during treatment. In blue, before or after treatment.s

### Illumina resistome data for surveillance purposes

To evaluate the use of MetaPointFinder for surveillance purposes, we selected and downloaded all Illumina read files from a study by Munk et al [12] describing the use of metagenomics for AMR surveillance of sewage from 101 countries from all continents (Figure 4). Sequencing data were processed using MetaPointFinder and the resistant percentage of RDRs and relative abundance of resistant RDR were clustered and displayed as heatmaps. Similar to Munk et al, clustering on RDR data by continent is observed and some resistant RDRs are mainly found in Europe (Figure 4AB). Completely analysing the Munk et al dataset is outside the scope of this manuscript, but for instance the fraction of resistant pleuromutilin RDR ABC−F_type_ribosomal_protection_protein_Eat(A) is much higher in Europe than in Asia, LAC or Africa which is visible in the percentage analysis. High pleuromutilin resistance has been noted before in Europe because of the use of tiamulin in farm animals [28].

**Figure 4.**
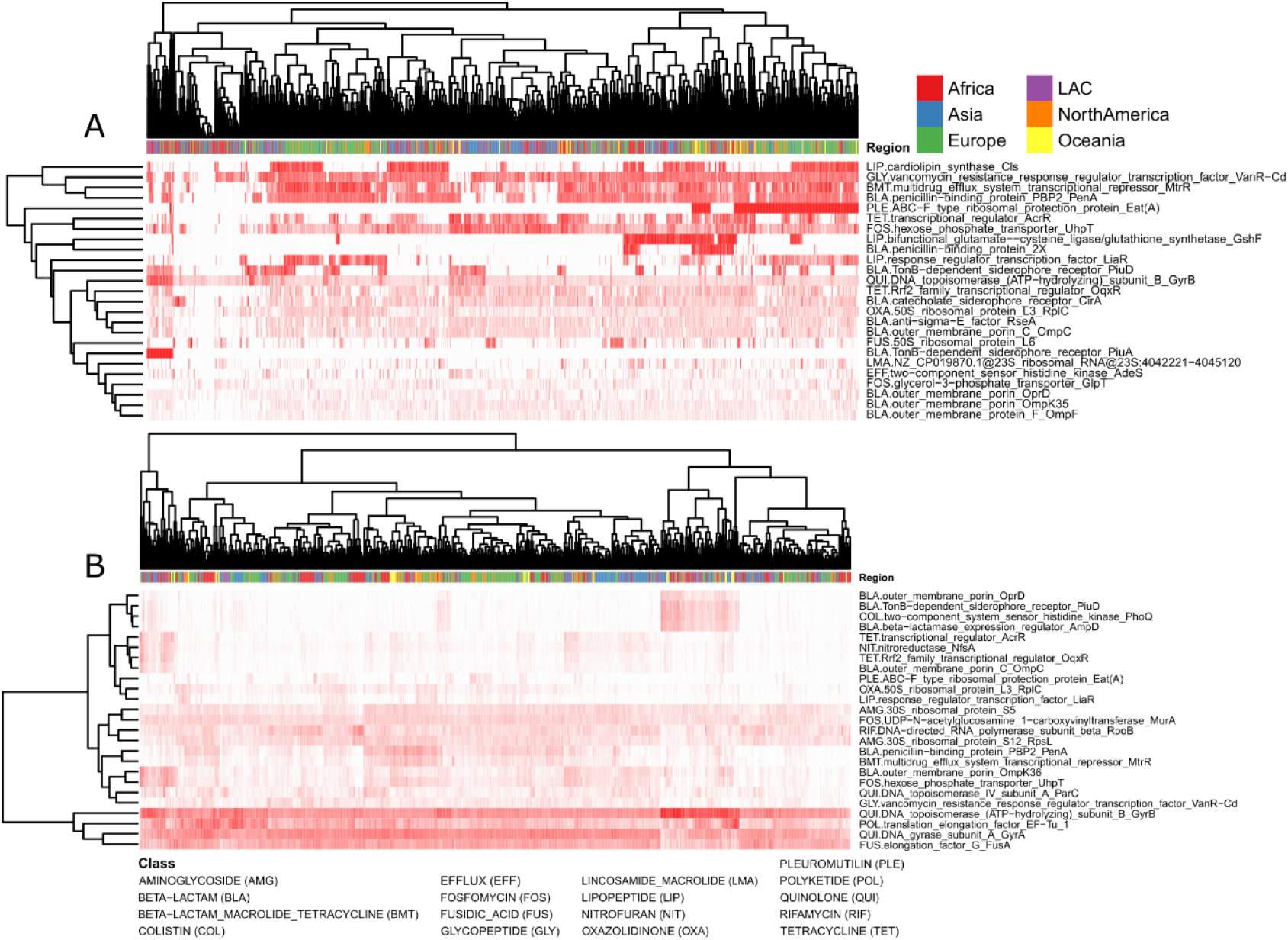
Heatmap of resistance of RDRs in sewage data from 101 countries. Clustered ARM heatmap showing the percentage resistance (A) and relative abundance (B) of the 25 most variable ARMs in the dataset from Munk et al [12]. LAC: Latin America and the Caribbean.

### Comparison of ARGs and ARMs in Nanopore sequencing data

No long-read benchmark metagenomics sequencing dataset is available with known quantities of resistance based on phenotypic antimicrobial resistance data. Based on the assumption that resistance caused by both ARGs and ARMs are a result of selection caused by exposure to antibiotics, we could assume that samples with high counts of ARGs may also have a high count of ARMs. To evaluate the performance of MetaPointFinder on Nanopore data, we downloaded all long-read metagenomes from the SRA with more than 1 gigabase per sample (n=1228) and we determined ARG counts per class by aligning all reads with the AMRFinderPlus protein database. The ARM counts were determined using MetaPointFinder with a sequence identity cutoff 70% to accommodate the higher expected error rates of long-read metagenomic datasets. The ARG and ARM readcounts were corrected for total read number. Spearman Rank and log+1 transformed Pearson correlation between ARG and ARM per sample was determined per class (Supplemental Data 1, Table 1). For many classes, a correlation was observed between ARG and ARM abundance. For colistin and macrolide drug classes, the observed correlation was low or absent, possibly because the number of samples with more than 10 reads with ARGs was low for colistin (35 samples) and the number of samples with ARM reads was low for macrolides (79 samples).

**Table 1.** Pearson and Spearman Rank Correlation of ARG and ARM abundance corrected for read depth.

| Class | Pearson | Pearson* | Log(x+1)<br>Pearson | Spearman | Spearman* |
| --- | --- | --- | --- | --- | --- |
| aminoglycoside | 0.14 | 0.19 | 0.32 | 0.32 | 0.33 |
| beta-lactam | 0.44 | 0.46 | 0.48 | 0.43 | 0.43 |
| colistin | 0.02 | -0.01 | 0.19 | 0.21 | 0.42 |
| fosfomycin | 0.25 | 0.38 | 0.64 | 0.57 | 0.65 |
| fusidic acid | 0.42 | 0.52 | 0.51 | 0.34 | 0.60 |
| macrolide | 0.00 | 0.04 | 0.18 | 0.23 | 0.10 |
| quinolone | 0.17 | 0.37 | 0.4 | 0.33 | 0.61 |
| rifamycin | 0.13 | 0.26 | 0.33 | 0.28 | 0.49 |
| tetracycline | 0.17 | 0.28 | 0.13 | 0.13 | 0.10 |

\* Samples with zero counts have been excluded. All r values above 0.056 are significant.

### Analysis of Nanopore deep sequencing data

In Bloemen et al. [29], a study to evaluate the ability of long-read sequencing to detect very low numbers of fluoroquinolone point mutations in deep sequenced data using haplotype-aware assembly methods, phasing of gyrase mutations from the metagenomics assemblies was challenging. Here, MetaPointFinder detected RDR substitutions directly in long reads. Both resistant and wildtype reads were detected, albeit lower in Sample A where detection completely failed in the original study (Table 2).

**Table 2.** Fluoroquinolone RDR counts of two deep metagenomic samples sequenced. WT=wildtype, RES=resistant.

| RDR | Sample A |  | Sample B |  |
| --- | --- | --- | --- | --- |
|  | WT | RES | WT | RES |
| DNA gyrase subunit A GyrA | 72 | 4 | 430 | 28 |
| DNA topoisomerase (ATP-hydrolyzing) subunit B GyrB | 138 | 2 | 148 | 4 |
| DNA topoisomerase IV subunit A ParC | 98 | 8 | 222 | 52 |
| DNA topoisomerase IV subunit B ParE | 104 | 8 | 126 | 2 |

### Amplicon datasets

Applying MetaPointFinder to a *gyrA*/*parC* amplicon sequencing study by Johnning et al [25], examining fluoroquinolone resistance in Swedish and Indian river water samples taken from different distances from a WWTP, revealed location-specific differences in *gyrA*/*parC* mutation burdens that were obscured by discarding a large fraction of reads in the workflow used by the authors of the original study (Table 4). Twice as many reads are mapped by MetaPointFinder than the original study’s custom workflow and a clear difference was observed in upstream samples (80 % vs 63%, Table 3). Swedish and Indian river water samples up- and downstream of a waste-water treatment plant contain different levels of ARMs, although statistical testing cannot be performed as there are no replicates. Surprisingly, the amplicon reads also encode hits to GyrB and ParE, likely because of the conservation of the genes encoding these proteins results in off-target amplification and subsequent identification by MetaPointFinder.

**Table 3.** Reads mapped to quinolone RDRs and percentage of RDRs resistant of total wildtype and resistant RDRs.

| Sample location with regards to WWTP | Total Read count | reads mapped Johnning et al | reads mapped MetaPointFinder | GyrA %resistant | GyrB %resistant | ParC %resistant | ParE %resistant |
| --- | --- | --- | --- | --- | --- | --- | --- |
| Indian 1.9 km upstream | 31384 | 3762 | 18893 | 80% | 1% | 4% | 5% |
| Indian 2.3 km downstream | 30094 | 2939 | 14644 | 75% | 1% | 65% | 6% |
| Indian 2.7 Km downstream | 27483 | 4215 | 12695 | 62% | 2% | 38% | 2% |
| Indian 17.5 km downstream | 26277 | 2262 | 12180 | 57% | 2% | 0% | 2% |
| Swedish 5-100 m upstream | 29010 | 4885 | 16412 | 63% | 4% | 23% | 3% |
| Swedish 25-230 m downstream | 26035 | 2719 | 13682 | 59% | 6% | 13% | 3% |

## Discussion

MetaPointFinder addresses a key gap in metagenomic resistome profiling, the identification of mutation-driven AMR, by directly querying known resistance-conferring substitutions at the read level. Compared with prior mutation mappers such as Mumame, our translated amino-acid and DNA alignment strategy improves recovery and quantification, especially for long reads or sequences with higher divergence from the reference sequences. Mumame relies on the free 32 bit version of USEARCH which is limited to files with a maximum of 4 gbytes [30], thereby limiting its use cases. We show that MetaPointFinder detects twice as much RDRs with 10-1000 fold less false positives as compared to Mumame, depending on read length and error rate.

While AMRFinderPlus and RGI catalogue known SNPs effectively on assembled isolate genomes, and RGI command-line can process metagenomes, these tools are not primarily designed for per-read ARM quantification of metagenomic data as species have to be selected *a priori*. Secondly, the difficulty with assembled metagenomes is that minor variant ARMs are lost in standard assemblies. In Bloemen et al [29], an attempt was made to use haplotype-aware assembly methods on two metagenomic samples, sequenced at a high depth, to phase out genomes from the same species, but this resulted in limited success. The authors only detected the gyrase mutation in one sample. In many cases, read-based analysis, like commonly used for ARGs, is the preferred method for ARMs as well as it allows for the identification of ARMs in metagenomes that might be variably present in different members of the same species. It does not rely on assembly, which requires higher read depths of both wildtype and resistant RDR as compared to read-based analysis.

Because MetaPointFinder not only scores resistant RDRs but also explicitly quantifies wild-type RDRs, the resistant proportion can be determined, similar to the proportion-based indicator-organism methods used in phenotypic surveillance; in both approaches, resistance is expressed as a fraction of the total population, resistant reads in metagenomes or resistant colonies in culture, reflecting the same underlying surveillance philosophy described by e.g. Dorado-García et al and EFSA/ECDC guidelines [31, 32]. In addition, MetaPointFinder can express mutation-driven resistance in terms of relative abundance within the metagenomic read pool, analogous to how ARG abundances are routinely quantified in resistome profiling. This relative abundance reflects the load of resistance and is important for risk assessment, as higher abundance is directly associated with increased risk of resistance dissemination [33–35].

Limitations of MetaPointFinder include reliance on known mutations and a single database source (AMRFinderPlus). Future improvements could include including more known ARM data from published sources. Another improvement could be the assessment of mutations from multiple database hits for a single RDR, thereby increasing sensitivity, possibly at the cost of specificity. A further improvement would be to include an extended, non-evidence based experimental database by also scoring conservative alternative substitutions informed by substitution matrices and probabilistic scoring, or machine learning-based predicting of similar mutations, or protein structure informed alternative substitutions. However, these are computer predictions that need to be used with caution until validated *in vitro*. We plan to release a predicted resistance mutation database addressing these potential improvements. A final limitation to using read based analysis is that resistances that require a combination of mutations in several genes, such as penicillin resistance in *S. pneumoniae* [36] cannot be evaluated as the presence of the combination of mutations in a single phased out genome sequence from a metagenome cannot yet be reliably determined.

## Conclusion

MetaPointFinder enables robust detection and quantification of known resistance-conferring point mutations directly from metagenomic reads, complementing gene-centric analyses and improving surveillance of mutation-driven AMR, especially for classes of resistance that are mostly driven by ARMs instead of ARGs.

## Data and Code Availability

MetaPointFinder repository https://github.com/aldertzomer/metapointfinder. Benchmark scripts: https://github.com/aldertzomer/metapointfinder/benchmark. Online version https://klif.uu.nl/metapointfinder/. Supplemental data 1 used to generate tables in figures in this manuscript: https://doi.org/10.5281/zenodo.17737473

## Acknowledgments

AZ and SN acknowledge funding from the Netherlands Organisation for Health Research and Development (ZonMw) [Grant 10430022010002]. AEM acknowledges funding from the UK Research and Innovation (UKRI) Medical Research Council (MRC) [Grant MR/Y034457/1], AEM and CC acknowledge funding from the Biotechnology and Biological Sciences Research Council (BBSRC) Institute Strategic Programme Microbes and Food Safety BB/X011011/1 and its constituent project BBS/E/QU/230002C (Theme 3, Flexible capabilities to reduce food safety threats and respond to national needs). The authors acknowledge support from the Joint Programming Initiative on Antimicrobial Resistance (JPIAMR) through the MEGAISurv project, and the BBSRC Europe Partnering Award BB/V01823X/1. The authors also acknowledge the use of ChatGPT (version 5; OpenAI) for assistance in code curation and grammar correction.

